# Slow rTMS to the left DLPFC enhances verbal memory formation

**DOI:** 10.1101/2021.03.02.433519

**Authors:** Mircea van der Plas, Verena Braun, Benjamin Johannes Stauch, Simon Hanslmayr

**Author notes:** Corresponding authors: Simon Hanslmayr. These authors contributed equally to this work (Mircea van der Plas, Verena Braun).

## Abstract

Encoding of episodic memories relies on stimulus-specific information processing and involves the left prefrontal cortex. We here present an incidental finding from a simultaneous EEG-TMS experiment as well as a replication of this unexpected effect. Our results reveal that stimulating the left dorsolateral prefrontal cortex (DLPFC) with slow repetitive transcranial magnetic stimulation (rTMS) leads to enhanced word memory performance. 40 healthy human participants engaged in a list learning paradigm. Half of the subjects (N=20) received 1 Hz rTMS to the left DLPFC while the other half (N=20) received 1 Hz rTMS to the vertex and served as a control group. Subjects receiving left DLPFC stimulation demonstrated enhanced memory performance compared to the control group. This effect was replicated in a double-blind within-subjects experiment where 24 participants received 1 Hz rTMS to the left DLPFC and vertex. In this second experiment, DLPFC stimulation also induced better memory performance compared to vertex stimulation. In addition to these behavioural effects, we found that 1 Hz rTMS to DLPFC induced stronger beta power modulation in posterior areas, a state which is known to be beneficial for memory encoding. Further analysis indicated, that beta modulations did not have an oscillatory origin. Instead, the observed beta modulations were a result of a spectral tilt, suggesting inhibition of these parietal regions. These results show that applying 1 Hz rTMS to DLPFC, an area involved in episodic memory formation, improves memory performance via modulating neural activity in parietal regions.

## Introduction

We are able to encode and store episodes that are rich in detail, filled with information and highly associative [1]. The first crucial step in forming episodic memories consists of processing the information at hand [2]. Before an event can be stored for later access it has to be represented [3]. This involves posterior neocortical areas processing different sensory inputs under top-down control of prefrontal regions [4,5]. Being able to enhance this process via brain stimulation could prove invaluable not only for therapeutic interventions but also for gaining knowledge about how our brain accomplishes the complex task of forming episodic memories.

The left dorsolateral prefrontal cortex (DLPFC) has been demonstrated to play a role in memory formation (for a review, see [6]). Stimulation at the DLPFC during encoding has been shown to reduce performance on verbal episodic memory tasks [7,8]. These reductions in performance have been mainly achieved with facilitative stimulation protocols (20 Hz stimulation). Thus, it seems that left DLPFC activity might have an inverse relationship to memory performance. Thereby, by inhibiting the left DLPFC, one would expect to see an increase in memory performance. Slow repetitive Transcranial Magnetic Stimulation (rTMS) has been shown to have an inhibitory effect on cortical areas [9–12].

Monitoring the ongoing electrophysiological activity, with electroencephalography (EEG) can inform the mechanisms that lead to a given behavioural observation. We were particularly interested in monitoring the ongoing spectral profile, oscillations in the alpha-beta frequency band typically show a reduction in power during successful memory processing (see [13] for a review), which might reflect more efficient stimulus processing [3,15].

We here report an incidental finding from the dataset of an existing study [15] in which the authors examined the role of the left DLPFC in voluntary forgetting. We re-analysed their rTMS-EEG dataset and found that 1 Hz rTMS applied to the left DLPFC during encoding of verbal material enhances memory performance. We further found that this rTMS-induced enhancement of memory performance co-occurred with stronger beta-power decreases, a state which is known to be beneficial for stimulus processing [16]. To ensure that the memory enhancing effects of rTMS are replicable, we conducted a second experiment which confirmed the memory enhancing effect of left DLPFC stimulation (experiment 2).

## Results

### Experiment 1: Behaviour

Participants were presented with two lists of ten words per encoding-retrieval run. Following each list, they were instructed to remember (i.e. keep in mind) the list just presented. After undertaking a short distractor task, participants were asked to recall all words from the two word lists just presented. The experimental group received 1 Hz rTMS to the left DLPFC during encoding of the second list and the control group received stimulation to the vertex (see Figure 1).

**Figure 1.**
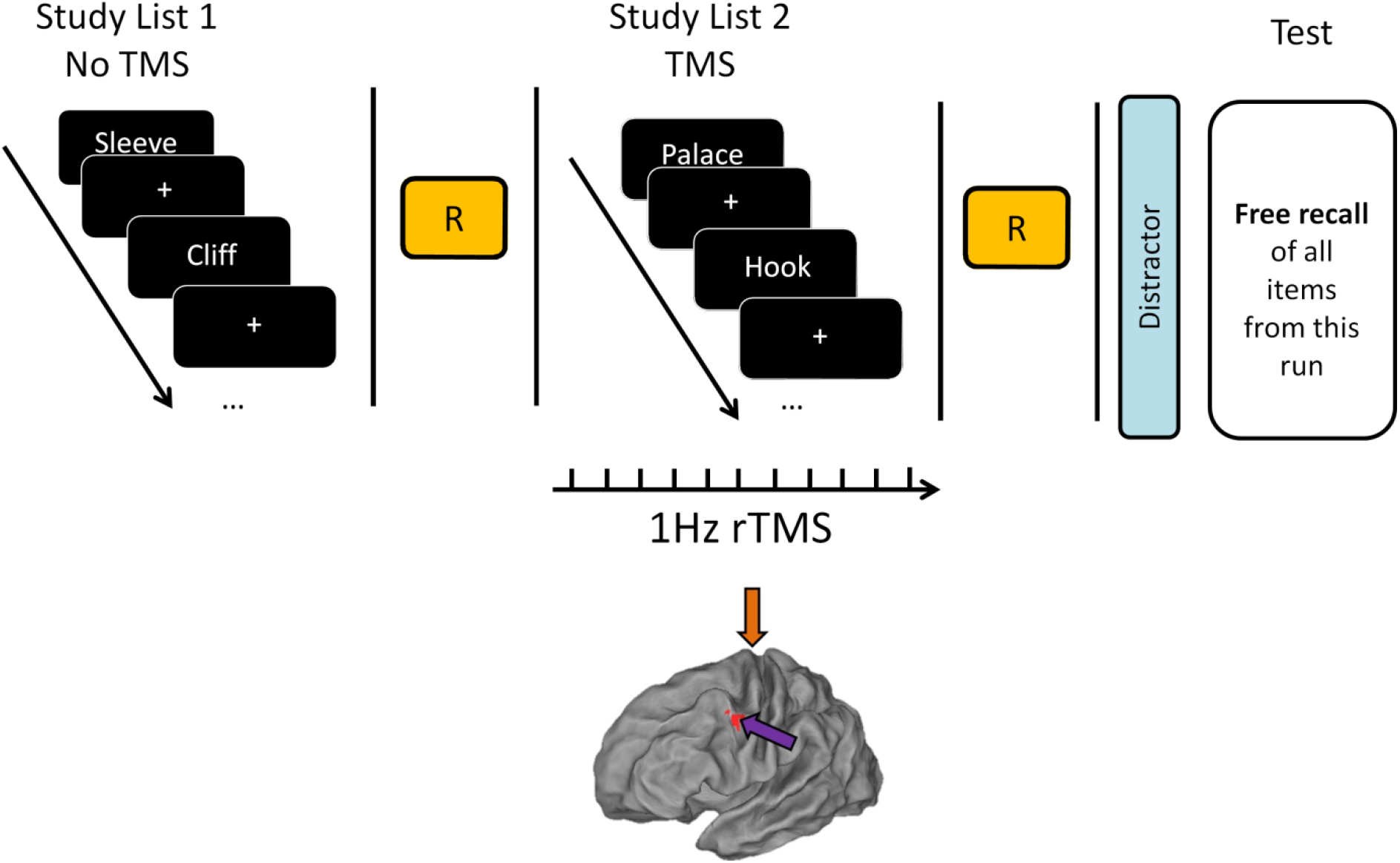
Experimental design. Arrows indicate stimulation site (DLPFC=purple, vertex=orange). Participants were asked to study two lists of 10 words. During encoding of list 2, 45 pulses of 1Hz rTMS were applied to the left DLPFC (MNI coordinates: −45, 6, 39) or vertex. Memory performance was assessed as percentage of correctly recalled words per list.

#### Behaviour

To test the effect of rTMS on memory performance we conducted a 2 (List 1 vs List 2) x 2 (DLPFC vs vertex) mixed ANOVA. There was a significant positive effect of DLPFC stimulation on memory performance (main effect rTMS, F(1,38)=5.096, p=0.03, η^2^_p_=0.118) and a significant difference between memory for the first and second list (main effect list, F(1,38)=17.242, p<0.001, η^2^_p_=0.312). We also found a significant rTMS x LIST interaction (F(1,38)=8.837, p=0.005, η^2^_p_=0.189). Post-hoc independent samples t-tests revealed that the DLPFC group showed better memory performance compared to the vertex group for words presented during rTMS application (list 2, t(38)=2.820, p=0.008, Cohen’s d=0.892 Figure 2D), but not for words presented before rTMS application (list 1, t(38)=1.399, p=0.170, Cohen’s d=0.443, Figure 2B). Hence, the effects were specific to the application of rTMS to the left DLPFC.

**Figure 2:**
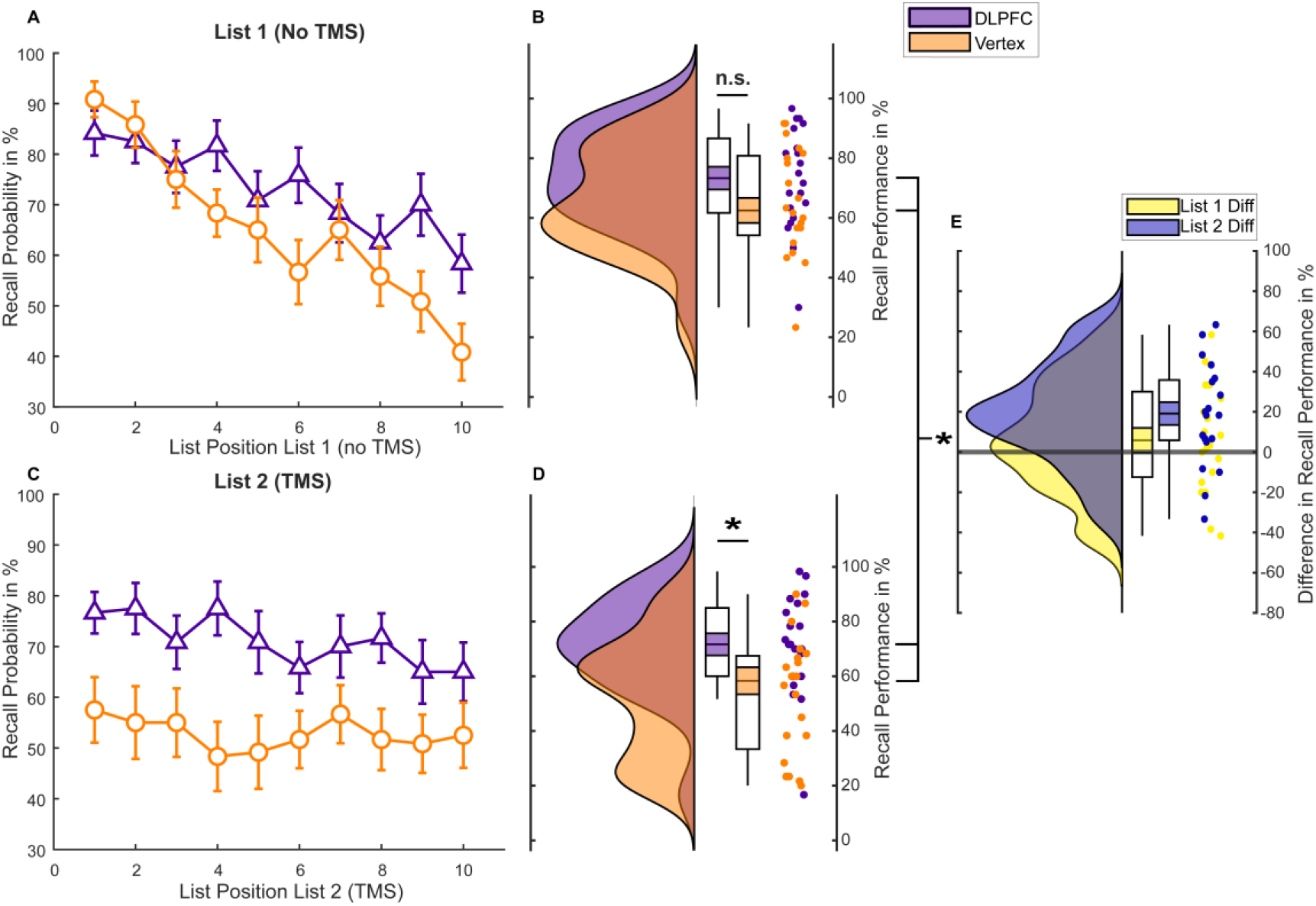
Memory performance experiment 1. A) Serial position curve for List 1 words. Error bars depict standard errors of the mean. B) Raincloud plots of average memory performance for List 1 words across all blocks with paired boxplots [17]. Coloured area within the box-plots indicate the standard error, while the circles depict individual data points. C) Serial position curve for List 2 words. Error bars depict standard errors of the mean. D) Memory performance for List 2 words. E) Difference in average memory performance between the DLPFC and vertex condition for each list (List 2 = Stimulation).

In an exploratory follow-up ANOVA we investigated a possible effect of rTMS on serial position to assess whether left DLPFC stimulation affected the likelihood of recalling a word as a function of its list position [18]. Analysis of serial position curves revealed a significant LIST x POSITION x rTMS interaction (F(9,342)=2.435, p=0.011, η^2^_p_=0.06). To unpack this 3-way ANOVA, we calculated two 2-way ANOVAs for each list separately. These ANOVAs showed a significant POSITION x rTMS interaction for list 1 (F(9,342)=2.703, p=0.005, η^2^_p_=0.066), but no significant POSITION x rTMS interaction for list 2 (F(9,342)=0.893,p=0.532, η^2^_p_=0.023, Figure 2C). The significant interaction in list 1 was due to enhanced recall rates for late position words in the DLPFC group compared to the vertex group (see Figure 2A). These results suggest that online rTMS to the left DLPFC equally increased memory performance in list 2 regardless of position, whereas for list 1 only late position words benefitted from stimulation.

### Experiment 1: EEG

Post-stimulus beta power decreases have repeatedly been associated with successful memory formation [13,19,20]. Therefore, we first tested whether the DLPFC group would show stronger post-stimulus (0 to 1 s) beta power decreases (13-30 Hz) for words that were later remembered (hits) compared to the vertex group for list 2 trials. In order to test for a difference in this time and frequency window of interest the data were subjected to a cluster-based permutation test [21]. The results show significantly stronger beta power decreases (13-30 Hz) post-stimulus during DLPFC stimulation compared to vertex stimulation. This effect was evident over bilateral posterior sites post-stimulus (p_corr_<0.05, Figure 3B; right post-stimulus topography). No effects were obtained for alpha (8-12Hz) or theta (4-7Hz) frequency bands in this time window. The time frequency plot at this the negative electrode cluster, as well as the time course of beta power, is shown in Figure 3A and 3C. Beta power showed a clear modulation due to rTMS with regards to word onset in the posterior electrode cluster. Specifically, stronger beta power pre-stimulus and lower beta power post-stimulus was observed during DLPFC stimulation compared to vertex stimulation.

**Figure 3.**
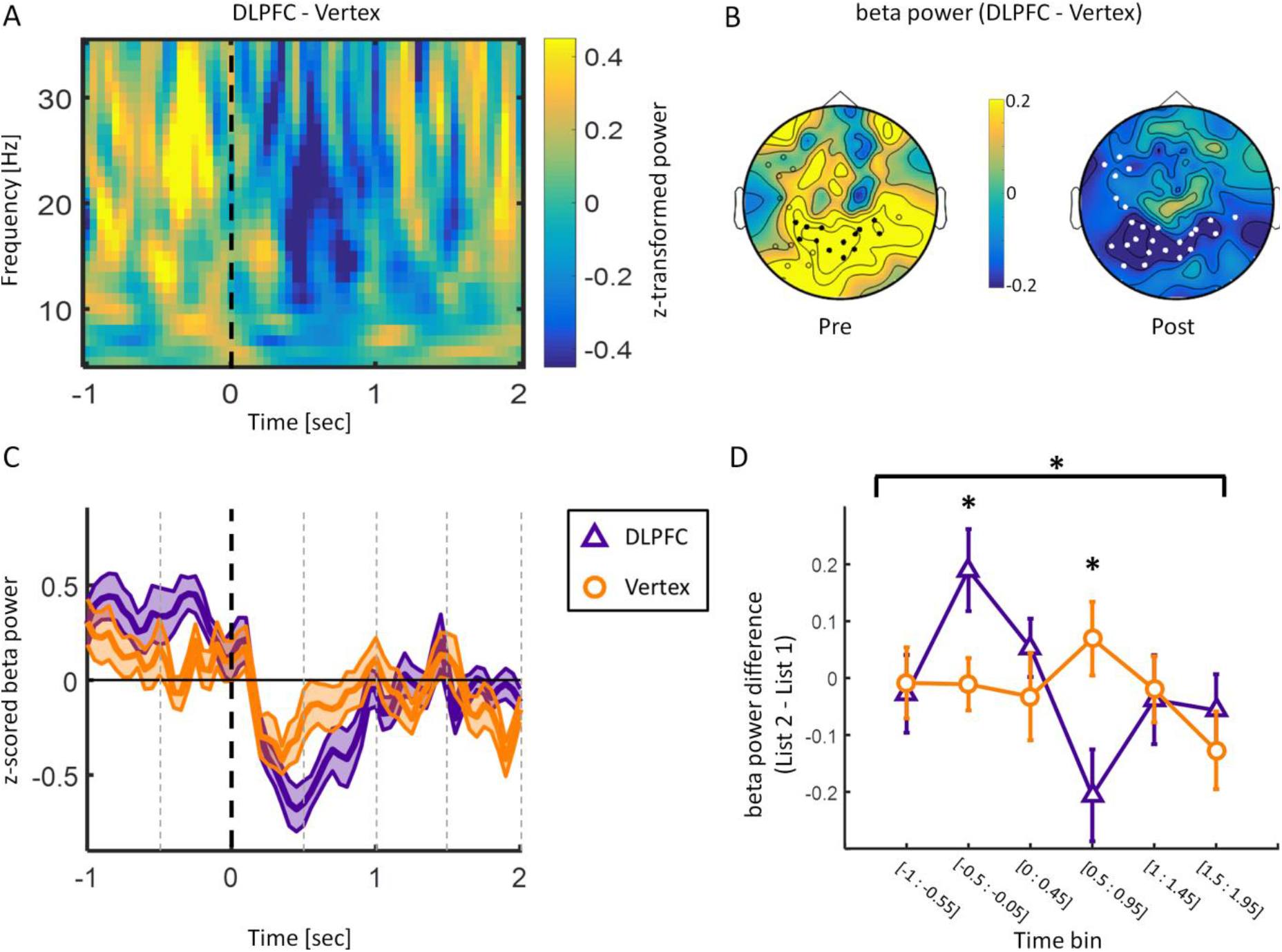
EEG results (only later remembered trials analysed). A. Time frequency plot for the difference between DLPFC and vertex during List 2 encoding averaged over electrode cluster demonstrating a significant negative difference (i.e. less power for DLPFC compared to vertex) between the DLPFC and vertex group in the beta frequency range post stimulus. Dashed line indicates word onset. B. Topographies depicting beta power (13-30 Hz) difference between DLPFC and vertex stimulation in time windows of interest (pre: −0.5 s to −0.05 s; post = 0 to 1 s). White circles depict significant negative electrode cluster post-stimulus. Black circles show electrodes within the negative cluster showing a positive difference pre-stimulus. C. Time course of beta power (13-30 Hz) averaged over the negative electrode cluster shown in B. Shaded area represents standard error of the mean. Black dashed line indicates word onset. Grey dashed lines depict time bins. D. Beta power difference (List 2 - List 1) over significant negative electrode cluster split by rTMS. Error bars show standard error of the mean. Data was split into six non-overlapping time bins: [-1s to −0.55s]; [−0.5s to −0.05s]; [0s to 0.45s]; [0.5s to 0.95s]; [1s to 1.45s]; [1.5s to 1.95s].

We further explored this beta power modulation to investigate whether it was specific to stimulation trials. Data from −1 s to 1.95 s relative to stimulus onset were split into six non-overlapping time bins (see Figure 3D) for List 1 and List 2 trials for the DLPFC and vertex group respectively. Data averaged over the significant negative electrode cluster were then subjected to a TIME (time bins) x LIST (List 1 vs List 2) x GROUP (DLPFC vs vertex) ANOVA which revealed a significant LIST x TIME x GROUP interaction (F(5,190)=2.707, p=0.022, η^2^_p_=0.066). Post-hoc independent samples t-tests revealed significant increases in beta power pre-stimulus (−0.5 s to-0.05 s: t(32.347)=2.384, p=0.023, Cohen’s d= 0.754) and decreases in beta power post-stimulus (0.5 s to 0.95 s: t(38)=-2.678, p=0.011, Cohen’s d=-0.847) in the DLPFC group compared to the vertex group (Figure 3D). These results indicate that 1 Hz rTMS at DLPFC modulated beta power predominantly in trials where the stimulation was applied.

### Experiment 1: Spectral Tilt vs Oscillations

Recent research suggests that some broadband memory-related effects are driven by a shift in spectral tilt (i.e. aperiodic components) rather than a change in narrow band oscillations (i.e. periodic components) [22]. To investigate if the above reported effect of DLPFC stimulation on beta power is due to a change in oscillatory activity or a change in spectral tilt, we separated power-spectra into periodic and aperiodic components using the FOOOF toolbox [23]. We performed a 2 (pre vs post: TIME) x 2 (DLPFC vs vertex: STIMULATION) repeated-measurements ANOVA on the periodic and aperiodic components respectively, with TIME as a within subjects factor and STIMULATION as a between subjects factor. We observed a significant interaction effect for the aperiodic component, as reflected by the exponent and offset of the aperiodic component: Exponent: PREPOST x STIMULATION: F(1,38)=5.900, p=0.020, η^2^_p_=0.134; Offset: PREPOST x STIMULATION F(1,38)=5.646, p=0.023, η^2^_p_=0.129 (see Figure 5, for the distributions of the separate components see supplementary figure S1). No such interaction effect was observed for the ANOVA investigating the periodic/oscillatory activity in the beta frequency band (PREPOST x STIMULATION: F(1,27)=0.652 p=0.426, η^2^_p_=0.024). These results suggest that the time-frequency representation was mainly driven by the aperiodic component, rather than narrow band oscillatory beta activity. In particular, the results suggest that DLPFC stimulation leads to a steeper aperiodic component where power decreases more quickly as frequency increases.

### Experiment 2: Behavioural Replication

Experiment 1 revealed that 1Hz rTMS to the left DLPFC can increase memory performance for words that were presented during the stimulation compared to a control group. Enhancing long-term memory through rTMS would indeed be an important finding; especially with such a low frequency stimulation technique that does not require intracranial electrical stimulation or lengthy protocols. Given that our behavioural results were an incidental finding, we attempted an internal replication of the behavioural effect. To rule out any unspecific differences between the groups which might have contributed to the effects, we changed the study design to a within-subjects experiment. Furthermore, in this experiment the participants as well as the experimenter who interacted with them and scored their memory performance were naïve to the predicted effects of left DLPFC stimulation on memory. Other results of this study have already been reported [24].

To test whether DLPFC stimulation leads to enhanced recall rates compared to vertex stimulation, we conducted a 2 (List 1 vs List 2) x 2 (DLPFC vs vertex) repeated-measurements ANOVA. We found a significant main effect for stimulation in the 2×2 rm-ANOVA, showing that DLPFC stimulation indeed led to higher memory performance compared to vertex stimulation (main effect rTMS, F(1,22)=6.778, p=0.016, η^2^_p_=0.236). We did not, however, observe a significant effect for list or a significant interaction (main effect List, F(1,22)=2.943, p=0.100, η^2^_p_=0.118; interaction Effect List X rTMS, F(1,22)=0.009, p=0.926, η^2^_p_<0.01). Post-hoc t-tests revealed a significant difference in recall performance between the DLPFC compared to the vertex condition for list 2 words, during the actual stimulation (t(22) = 2.38, p = 0.026, Cohen’s d=0.496; see Figure 5D). This comparison was not statistically significant for list 1 words (t(22) = 1.754, p = 0.093, Cohen’s d=0.366; see Figure 5B). This pattern suggests, that left DLPFC stimulation, once again led to enhanced memory performance compared to vertex stimulation. Analysis of the serial position curves (Figure 5A and C) revealed that recall performance across positions did not differ between the DLPFC and vertex condition in either of the two lists (rTMS x LIST x POSITION: F(9,198)=1.061, p=0.394, η^2^_p_=0.046; List 1: rTMS x POSITION F(9,198)=1.612, p=0.114, η^2^_p_=0.068; List 2: F(9,198)=0.811, p=0.607, η^2^_p_=0.036).

For most of the participants (N=19), the order in which words were recalled was also available. This allowed us to assess the amount of temporal clustering [25] for list 2 words (Procedure is explained in depth in [26]) and to examine whether DLPFC stimulation affected the amount of contextual error. Such an effect would be predicted by theories implicating the DLPFC in organizing memory material into temporal clusters [27]. A two-tailed dependent samples t-test was conducted to compare temporal clustering between DLPFC and vertex trials. There was no difference in contextual error between the DLPFC and vertex condition (list 2: t(18) = −0.231; p = 0.82, Cohen’s d=-0.053) indicating that the memory enhancement effect of left DLPFC stimulation cannot be attributed the increased temporal clustering of the words. Rather, DLPFC stimulation improved memory performance for each item independently.

Since experiment 1 and experiment 2 used virtually the same paradigm, we performed a continuously cumulative (weighted fixed-effect) meta-analysis over the two studies, in order to gain a more accurate estimate of the observed stimulation effect [28,29]. We found that stimulation on the left DLPFC significantly boosts memory performance for both List 1 and List 2 words across the two studies (g = 0.32 [0.01, 0.63]; g = 0.40 [0.15, 0.65]) (see Figure 6).

## Discussion

We demonstrated in two experiments that 1 Hz rTMS delivered to the left DLPFC during episodic memory encoding boosts memory performance. Participants encoded two lists of words and received 1 Hz rTMS during word presentation. In a subsequent free recall test, participants recalled significantly more words from lists in which they received left DLPFC stimulation compared to vertex stimulation. The accompanying serial position and contextual clustering analyses suggest that left DLPFC stimulation enhances stimulus processing at a word-specific level without affecting associations between words. Simultaneously recorded EEG data for the first experiment indicated that 1Hz rTMS to the left DLPFC strengthened event-related power decreases in the beta frequency band in posterior areas. This was represented by higher beta power before word onset and lower beta power after word onset in the DLPFC group compared to the vertex group. Taken together our results show that slow rTMS can enhance memory performance, and that this memory enhancement effect was associated with increased stimulus induced beta power decreases, an established correlate of memory function [13].

Power decreases in the alpha/beta frequency range is traditionally associated with stimulus processing in general [30]. While power increases in these frequency bands have been linked to inhibition of irrelevant or potentially interfering information, event-related power decreases (i.e. disinhibition) have been observed over areas actively involved in stimulus processing [31–33]. This beta power reduction has previously been shown to be vital for successful encoding of verbal material [34–36]. This makes sense conceptually, as areas in the MTL can only bind information that has been appropriately processed in down-stream neocortical areas [37]. Given its importance in information processing and representation, reduced activity in the alpha/beta frequency bands has been proposed to reflect active involvement of cortical areas during encoding of episodic memories [3,13]. Additionally, TMS has been shown to have network wide effects, which can extend throughout the brain [38,39]. Consequently, it appears that the DLPFC stimulation, somehow encourages stimulus processing in parietal and occipital areas, as reflected in the decreased power in those areas. However, a slightly different interpretation could be made considering the result of the analysis separating the periodic and aperiodic components. The observed power changes seem to result from an upwards (or clock-wise) rotation in the spectral tilt as observed by the increasing exponent and offset components, rather than a change in oscillatory components (see Figure 4). This type of rotation has been associated with increased inhibition [40] suggesting that the frontal stimulation has an inhibitory effect over the parietal cortex.

**Figure 4:**
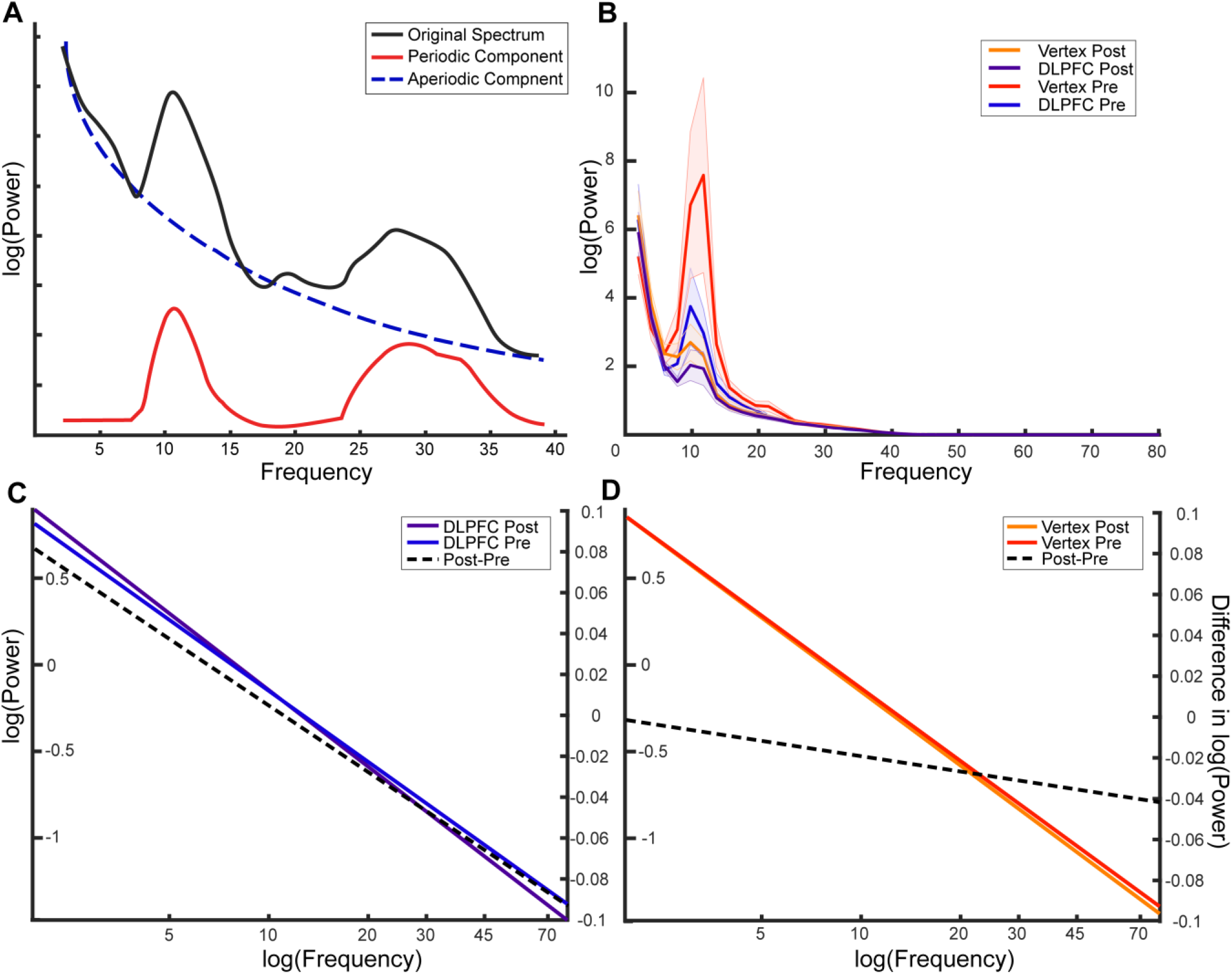
A) Schematic representation of the different components in a given power-spectrum. The black line represents a typical power-spectrum that is to be separated. The blue line is the corresponding log function following removal of the periodic peaks, thereby representing aperiodic properties of the signal. B) Power spectra separated by each condition. Shaded area indicates standard error. C)-D) Line plots of the mean aperiodic component before and after item presentation for the DLPFC and vertex Condition respectively. The right axis relates to the plotted post-pre difference (dotted line).

**Figure 5.**
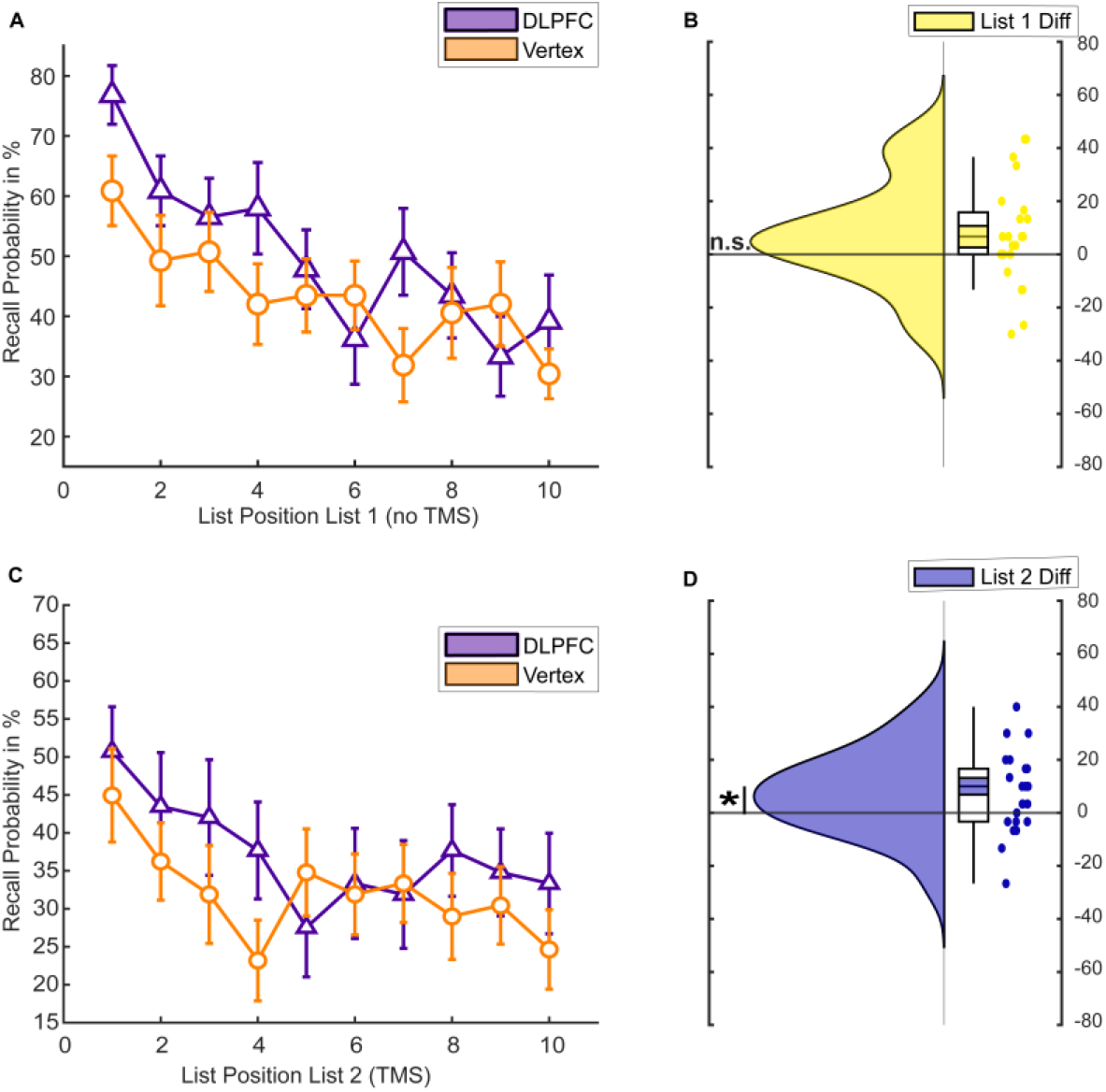
Memory performance experiment 2. A) Serial position curve for list 1 (N=23). B) Raincloud plots of memory performance for list 1 words (difference between DLPFC and vertex stimulation). Coloured area within the box-plots indicate the standard error, while the circles depict individual data points. C) Serial position curve for list 2. D) Raincloud plots of memory performance for list 2 words (difference between DLPFC and vertex stimulation).

**Figure 6:**
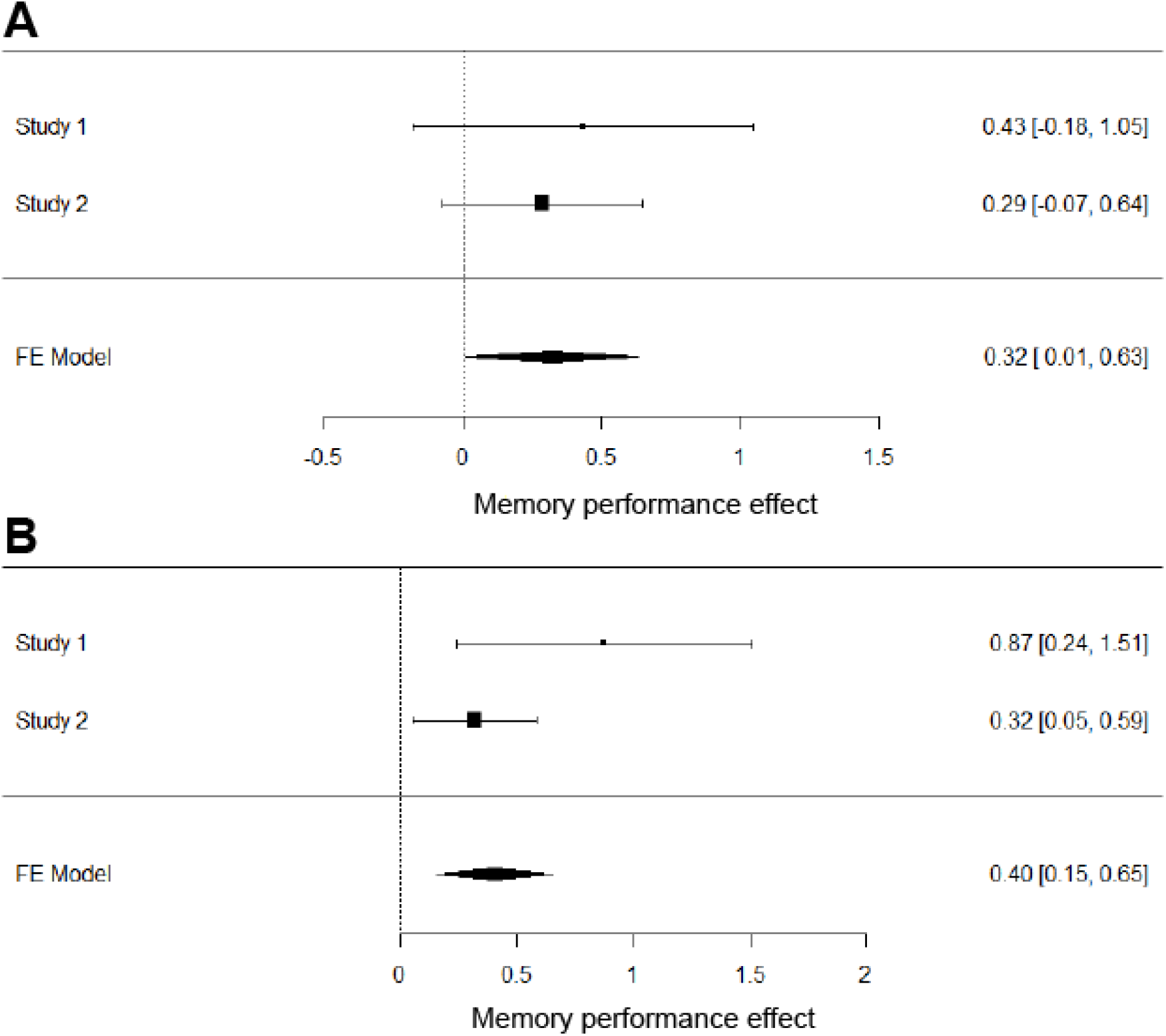
Forest plots of the meta-analytically combined DLPFC effect for List 1 (A) and List 2 (B) words. Error bars represent 95% confidence intervals, effect sizes were calculated as Hedge’s g.

This interpretation would be consistent with the fact that we used a stimulation protocol (1 Hz rTMS) that is usually considered to have inhibitory effects on cortical excitability [9,41]. Such an interpretation would be consistent with fMRI studies showing that decreased activity in ventral parietal regions are usually positively correlated with memory encoding[42]. This interpretation would also be consistent with other studies reporting a reduction in memory performance when stimulating the left DLPFC with parameters considered to increase excitability (i.e. 20 Hz; [7,8]). The behavioural effects observed in the two experiments described here therefore suggest an inhibitory relationship between the left DLPFC and verbal memory encoding. Further, the EEG results suggest that inhibition of the left DLPFC boosts event-related beta power decreases in the service of memory formation. This latter finding suggests that the DLPFC might actively limit the amount of stimulus processing in this memory paradigm. Inhibition of the DLPFC consequently leads to disinhibition in parietal down-stream areas. Such reductions in parietal beta power have previously been associated with an increased capacity of information coded into the neural signal[43]. This increase in potentially coded information would then ultimately result in a better memory performance.

An important caveat of the above interpretation is that it rests on the assumption that online rTMS affects the brain in the same way as offline rTMS does. While rTMS is a method that has been around for decades, most of the mechanistic studies rely on offline effects, where stimulation is first applied and its effects on neural activity or task performance are measured afterwards. This is a consequence of the large artifact a TMS pulse induces in EEG and MRI measurements. Thus, it is conceivable that, while the offline 1Hz r TMS may have inhibitory aftereffects, these could result as type of rebound effect from the actual stimulation (and vice versa for the online 20 Hz stimulation employed in the other studies). There has also been a study that have called the inhibitory qualities of 1 Hz rTMS into question[44]. Moreover, the effects of TMS onto the wider network can differ quite drastically from the local effects [38]. Thus, one should not discount the possibility that the parietal decreases might not be a result of modulating the DLPFC activity per se, but rather might result from influencing the memory network as a whole, in which the DLPFC plays an important role. Another possible interpretation, that disregards possible facilitative or inhibitory effects of rTMS, is that given our remote effects during left DLPFC stimulation, 1Hz rTMS may have influenced the functional connectivity between frontal and posterior regions [45]. This enhanced connectivity would then lead to enhanced stimulus processing and improved memory performance as a result thereof.

Despite our robust behavioural results, care should be taken when interpreting behavioural rTMS effects. External effects arising from rTMS can influence behavioural measures even when an active control condition is used. DLPFC stimulation, for example, can lead to stronger muscle twitches and distraction than vertex stimulation [46]. This may be experienced as distracting and affect encoding performance accordingly. However, if this was the case, one would expect this to affect performance negatively rather than positively. Furthermore, several studies have found similar effects as those we report here using different stimulation techniques or stimulation in adjacent regions [47–49]. Additionally, Köhler and colleagues [50] showed that when participants received 7Hz rTMS to the left inferior prefrontal cortex during a semantic encoding task [50], their word memory performance was enhanced. Two control sites were additionally stimulated —the right inferior prefrontal cortex and a right parietal target. Only left prefrontal stimulation resulted in more high-confident hit rates. These findings strengthen our confidence that the results presented are not merely a by-product of unspecific side effects, such as muscle twitches.

Behaviourally the results in both experiments demonstrate a positive effect of left DLPFC stimulation on memory performance in general. However, the results of the two experiments also differed slightly. Considering the first experiment, the memory effect was not only specific to the DLPFC stimulation condition compared to the vertex condition, but also specific for list 2 words, i.e. those words that were presented during rTMS. This finding was not replicated in the second experiment where there was no significant interaction between word list and stimulation condition. A possible reason might be carry over effects between lists. However, if this was the case, then the List by Stimulation interaction should also be absent in the first study. The only difference between the two experiments was that Experiment 1 had a between-subject design, while Experiment 2 had a within-subject design. Conceptually, there is no reason why the two designs would affect the difference between lists, as carry-over effects should still be present when a subject is only exposed to the DLPFC stimulation condition without an accompanying vertex stimulation condition. The results of the meta-analysis do support the possibility, that the significant interaction in the first study might be a false positive, because it suggests increases in memory performance for both lists across the two studies, thereby suggesting that rTMS during the second list might also enhance memory for previously encoded, but unstimulated items.

Lastly, as we analyse data recorded in directed forgetting paradigms, it is unclear if our results generalize to other types of memory tasks. However, considering other work on DLPFC stimulation and episodic memory, the involvement of the DLPFC in episodic memory encoding in general seems to hold across tasks [6,7,8]. Future research could clarify this by stimulating the DLPFC with 1 Hz rTMS during more general episodic and relational memory tasks.

## Conclusion

Our results indicate that 1 Hz rTMS applied to the left DLPFC during encoding of verbal material can enhance memory performance. This effect was linked to a well-known physiological correlate of memory formation: beta power decreases. Given the need for replication studies in general [51] and for brain stimulation effects in particular [52], we set out to replicate the initial incidental finding. In order to control for inter-individual differences [53–55], we replicated our original result in a within-subjects investigation. The results of this second experiment replicated the memory enhancement effect resulting from 1 Hz left DLPFC stimulation. Therefore, online 1 Hz rTMS at left DLPFC appears to be an effective means of enhancing cognitive function in a memory task with potential applicability ranging from basic research to clinical intervention. Future studies should further explore how exactly 1Hz rTMS to the left DLPFC gives rise to more pronounced beta power decreases in posterior areas and enhanced memory as a result thereof.

## Material and Methods

### Experiment 1

#### Subjects

The data reported here was collected as part of a larger study (reported in [15] experiment 2). 48 healthy human participants were tested and subjects were randomly assigned to one of the two stimulation conditions. After artefact rejection and inspection of the EEG data, 40 participants remained in the sample, resulting in 20 participants per group (DLPFC group: mean age = 21.7, range 18-26, 8 males; vertex group: mean age = 22.3, range 18-27, 6 males). All participants were right handed, had normal or corrected-to-normal vision, reported no history of neurological disease or brain injury, and were screened for contraindications against rTMS [10]. Informed consent was acquired from each subject prior to the experiment. The study was approved by the ethics committee of the University of Konstanz.

#### Task and Stimulus Material

The stimulus material consisted of 240 nouns derived from the MRC Psycholinguistic Database [56]. The material was translated into German and divided into 24 lists of 10 words. The lists were matched according to word frequency, number of letters, number of syllables, concreteness, and imageability [15]. The presentation of the lists was counterbalanced across subjects. Each list was presented equally often across four conditions (Forget List 1, Forget List 2, Remember List 1, Remember List 2). The data was collected as part of a study that focussed on the causal involvement of the left DLPFC in voluntary forgetting (reported in [15], experiment 2). Participants performed 12 encoding-recall runs. In each run, particpants were presented with two lists of 10 words. After having studied the first 10 words, a cue was presented for 5 s, prompting participants to either forget the previously studied words or to continue remembering this list. The second list of 10 words was always followed by a remember cue. For this study, only the six remember runs, i.e. runs in which the first and second list had to be remembered, are included in the analysis. The words were presented in randomized order one at a time for 2.5 s, with a variable inter-stimulus interval of 1.5-2.5 s (during which a fixation cross was shown). After a short distractor task of 2 min (counting backwards in steps of 3 from a random number), participants were asked to freely recall as many words from this run as possible in any order. Participants’ reponses were recorded manually by the experimenter outside of the EEG room.

#### rTMS

During encoding of List 2, 45 pulses of 1Hz rTMS were applied at 90% resting motor threshold. One group of participants received rTMS to the left DLPFC, while the control group received rTMS to the vertex. There was no relationship between the timing of the rTMS pulses and the stimulus presentation. rTMS was delivered using a Magstim Rapid2 stimulator with a figure-of-eight air filmed cooled coil (magstim; www.magstim.com). Prior to the main experiment, individual T1-weighted MRI scans were acquired with a 1.5T Philips scanner. In order to assure that the exact regions of interest were targeted, the stimulation was guided by a neuronavigation system (ANT-Visor; www.ant-neuro.com). Individual MRI scans were co-registered with the position of the rTMS coil and the precise targeting of the stimulation sites was monitored throughout the experiment. The coil was approximately angled 45° from the midline axis of the participant's head with the handle pointing backwards and laterally. The MNI coordinates for DLPFC stimulation were x=-45, y= 6, z=39 [14].

#### EEG recording and preprocessing

EEG was recorded throughout the task from 128 electrodes in an equidistant montage (ANT; www.ant-neuro.com). Participants were seated in a shielded room and data were recorded with a DC amplifier (ANT) at a sampling rate of 2048 Hz; data were offline re-referenced to average reference. Individual electrode positions were digitized at the beginning of the experiment (Xsensor, ANT). EEG data were preprocessed and analysed using Fieldtrip [57]. Due to excessive artifacts in the EEG during rTMS [58], List 1 (no rTMS) and List 2 (during rTMS) trials were preprocessed separately. Preprocessing of rTMS-EEG data followed the guidelines and procedure outlined by Herring et al. [59] adapted to our dataset. EEG data were first cut into segments of −0.9s to 0.9s around the rTMS pulses. Data were visually inspected and data around the rTMS artifacts removed from further analysis. The epoched data were subjected to an independent component analysis (runICA). This allowed the removal of rTMS related artefact, eye-blink, eye movement and other remaining artefacts. The cleaned data epoched around word onset (−2s to 4s) were then downsampled to 500Hz. A low-pass filter (40 Hz cut-off) was applied and the data were visually inspected for remaining artefacts. Missing and rejected channels were interpolated (mastoids were removed resulting in 126 channels). For trials without rTMS (List 1), data were epoched −2s to 4s around the onset of the word, downsampled to 500 Hz, and low-pass filtered (40 Hz cut-off). After visually inspecting the data for artefacts, an ICA was applied in order to identify and remove ocular and muscle artefacts. The cleaned data were again visually inspected.

### Data Analysis

#### Behavioural Analysis

In order to assess the effect of stimulation on recall performance, a mixed ANOVA with the within subjects factor LIST (List 1 and List 2) and the between subjects factor rTMS (DLPFC and vertex) was performed. We further tested whether DLPFC stimulation influenced the likelihood of recalling words as a function on a words’ list position. To this end, serial position curves were calculated [18]. For every subject at every list position we coded whether a word was later recalled (1) or not (0). This was done for all six encoding-recall runs and subsequently averaged for every participant over the six runs. These data were then subjected to a 2 (DLPFC vs vertex) x 10 (position in list) x 2 (list 1 or list 2) ANOVA.

#### EEG Analysis

EEG data (−1.5s to 3 s) were subjected to a time-frequency decomposition (2 to 35 Hz in steps of 1 Hz) using Morlet wavelets (width 7) and z-transformed in order to enable analysis of post- as well as pre-stimulus activity [26]. As only negative clusters in the beta frequency range were expected, data from the DLPFC and vertex group were subjected to a one-tailed cluster based permutation test, averaged over beta (13-30 Hz) and the post stimulus time window of interest (0 to 1 s). Alpha values were set to 0.05. All further analyses were conducted on the electrode sites identified as showing significant differences in beta between the two conditions.

In order to further investigate the properties of observed power-changes, analyses were performed using the FOOOF toolbox [23]. This method uses simultaneous fitting of the aperiodic spectrum component as well as spectral peaks. For this we analysed the time-window of interest (resulting from the time frequency analysis) and a identically sized time-window before stimulus presentation. These components can then be analysed separately. We performed a 2 x 2 mixed repeated measure ANOVA (Pre vs Post (2) word presentation x DLPFC vs vertex (2) stimulation, for each component (the aperiodic exponent, the offset and the periodic peak power).

### Experiment 2

The data of experiment 2 was part of a larger study that focussed on replicating the effect of rTMS on directed forgetting and is reported elsewhere (see[60]).

#### Subjects

24 healthy human participants took part in this experiment (mean age = 19.04, range 18-28, 5 male). All participants were right handed, had normal or corrected-to-normal vision, reported no history of neurological disease or brain injury, and were screened against contraindications against rTMS [10]. Informed consent was acquired from each subject prior to the experiment and participants were fully debriefed at the end. The protocol was approved by the ethics committee of the University of Birmingham.

#### Task and Stimulus Material

In this study, the participants as well as the experimenter interacting with the subjects were blind towards the hypotheses.

240 nouns were derived from the MRC Psycholinguistic Database [56] and divided into 24 Lists of 10 words. As in experiment 1, the lists were matched according to word frequency, number of letters, number of syllables, concreteness, and imageability [15]. The presentation of the lists was counterbalanced across subjects so that each list was used equally often across eight conditions (DLPFC-Forget List 1, DLPFC-Forget List 2, DLPFC-Remember List 1, DLPFC-Remember List 2, vertex-Forget List 1, vertex - vertex List 2, vertex Remember List 1, vertex - Remember List 2). Participants performed 12 encoding-recall runs, split by stimulation condition. Whether the six DLPFC runs or the six vertex runs were conducted first was counterbalanced across subjects. The task was the same as in experiment 1. For this study, only the three remember runs per stimulation condition are included in the analysis. Participants’ responses were recorded manually inside the testing room.

#### rTMS

The same stimulation parameters were used as in experiment 1. However, in this experiment, participants received both DLPFC and vertex stimulation in a blocked manner. The stimulation was delivered using a Magstim Rapid stimulator with a figure-of-eight coil (magstim; www.magstim.com). Prior to the main experiment, individual T1-weighted MRI scans were acquired using a 3T Philips Achieva MRI scanner. In order to assure precise stimulation, individual MRI scans were co-registered with the position of the rTMS coil and the stimulation was guided by a neuronavigation system (Brainsight; Rogue Resolutions; https://www.rogue-resolutions.com). The coil was held in place manually and the precision of the stimulation was monitored throughout the experiment. The same MNI coordinates as in experiment 1 were used.

#### Meta-Analysis

In order to combine the effect of stimulation over the two studies, a cumulative meta-analysis of the stimulation effect for the list 1 and list 2 items was performed using the R-package metafor [29]. The analysis was performed by computing effect sizes (Hedge’s g) for the individual relevant t-tests (independent and dependent for study 1 and 2 respectively), , which were then used to run a weighted fixed-effect meta-analysis [27,57].

## Acknowledgments

This research was funded by Deutsche Forschungsgemeinschaft [Emmy Noether Programme Grant HA 5622/1-1] and the European Research Council [Consolidator Grant Agreement 647954] awarded to Simon Hanslmayr. We would like to thank Benjamin Griffiths for advice on temporal clustering analyses and Nora Oehler for help with data collection for experiment 1.

## Supplementary Material: Separated FOOOF Components per Condition

**Supplementary Figure 1:**
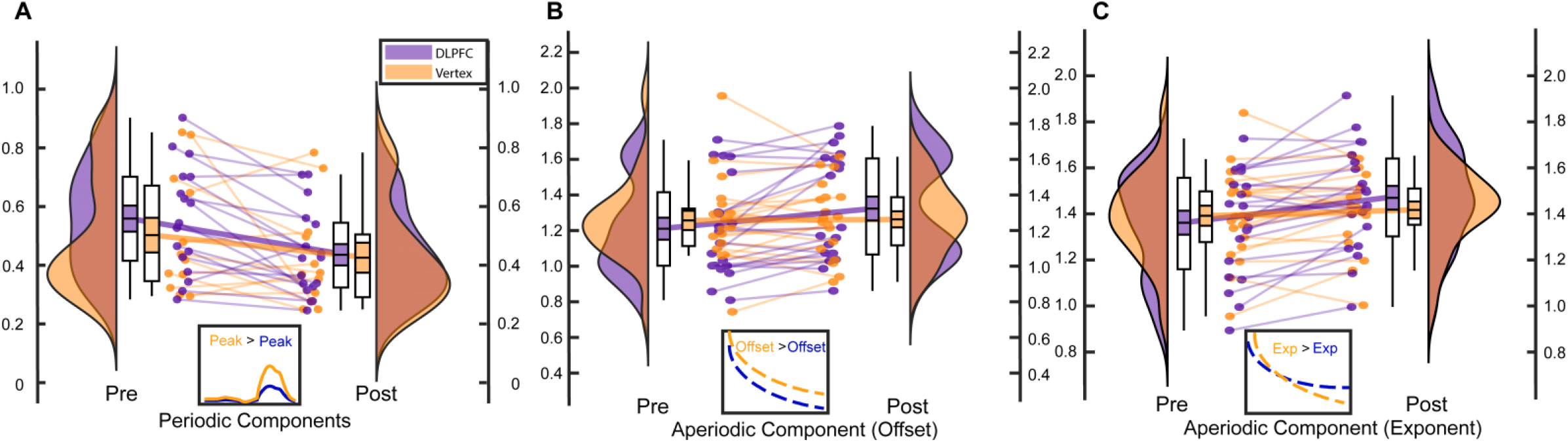
A) Raincloud plots of the beta periodic component for the DLPFC condition (purple) and the vertex condition (Orange), for the pre and the post stimulus condition respectively. B) Raincloud plots per stimulation condition for the offset of the aperiodical component at 0 Hz for pre and post word onset time windows. Yellow line represents an identical aperiodical component with an increased offset. C) Raincloud plots per stimulation condition for the Exponent of the aperiodical component Hz for pre and post word onset time windows. Yellow line represents an example for an identical component with a larger exponent. Each raincloud plot is paired with its respective boxplots. Coloured area within the box-plots indicate the standard error, while the circles depict individual data points for each participant respectively. The same subjects for the pre and post time-windows are connected by a line. The thick line illustrates the change in mean from pre to post.

## Supplementary Material: Addressing Difference in Trial Numbers

There was a considerable difference in the number of list 2 hits between the DLPFC and the vertex group because of the enhanced memory performance in the DLPFC group. (DLPFC: mean=23.1, SD=7.48; vertex: mean=17.25, SD=8.48). Power is not systematically biased by trial numbers, but we nevertheless tested whether this difference in trial numbers might have contributed to the observed effects. To this end, we randomly selected trials for each subject from the DLPFC group and matched these to the number of trials from subjects in the vertex group, ensuring that both groups have exactly the same trial numbers (mean: 17.25, SD: 8.48). As our main comparison of interest was the difference in beta power (13-30Hz) between the DLPFC and vertex group for list 2 trials, we conducted independent samples t-tests for data 0-1 s after word onset averaged over the negative electrode cluster identified earlier. This procedure was repeated 100 times, every time randomly selecting new subsets of trials for the DLPFC group. 100 t-tests on adjusted trial numbers revealed t values ranging from −3.9 to −2.377 (critical t for independent samples t-tests = 2.023; df=38). This analysis demonstrates that the difference in post-stimulus beta power decreases for list 2 words was not driven by differences in trial numbers.

